# Predator-induced shell plasticity in mussels hinders predation by drilling snails

**DOI:** 10.1101/124784

**Authors:** Zachary T. Sherker, Julius A. Ellrich, Ricardo A. Scrosati

## Abstract

Sessile invertebrate prey that detect waterborne predator cues often respond by strengthening their structural defenses. Experimental evidence of the functional significance of such modifications using field-raised organisms is lacking. This study addresses that gap using intertidal mussels and predatory dogwhelks from Atlantic Canada. During the spring and summer of 2016, we ran a field experiment that manipulated dogwhelk presence to test their nonconsumptive effects on mussel traits. Dogwhelk cues elicited thickening at the lip, centre, and base of mussel shells, although simultaneously limiting shell growth in length. As shell mass was unaffected by dogwhelk presence, a trade-off between shell thickening and elongation was revealed. Thickening was strongest at the thinnest parts of the shell. Using the field-raised organisms, a lab experiment found that dogwhelks took, on average, 55 % longer to drill and consume mussels previously exposed to dogwhelk cues than mussels grown without such a cue exposure. Dogwhelks drilled at the thinnest parts of the shell but, nonetheless, the consumed cue-exposed mussels had thicker shells at the borehole than the consumed mussels not exposed to cues, which likely explains the observed difference in handling time. As handling time normally decreases predation success, this study indicates that the plastic structural modifications in mussels triggered by dogwhelk cues in the field hinder predation by these drilling predators.

## INTRODUCTION

Nonconsumptive effects (NCEs) of predators on prey mediated by chemical cues are ubiquitous in aquatic systems (Ferrari et al. 2010, Brönmark & Hansson 2012). For example, when aquatic prey detect waterborne predator cues, short-term responses often include behavioural changes such as moving away or reducing feeding activities to decrease predation risk (Keppel & Scrosati 2004, Molis et al. 2011). Longer-term responses include the phenotypically plastic strengthening of morphological defenses, especially in prey with little or no escape capabilities, such as slow-moving and sessile species (Leonard et al. 1999, Nakaoka 2000, Freeman & Byers 2006). Predator NCEs may ultimately influence prey demography (Ellrich et al. 2015) and, indirectly, the abundance of other species in the community (Weissburg et al. 2014, Matassa et al. 2016). Thus, NCE research has become an important part of ecology. The present contribution investigates the functional significance of structural changes in sessile prey that are triggered by waterborne predator cues.

Morphological changes in shell-bearing invertebrate prey can be induced by waterborne cues from predatory snails, crabs, and sea stars (Reimer & Tedengren 1996, Smith & Jennings 2000, Cheung et al. 2004, Freeman 2007, Newell et al. 2007, Lord & Whitlatch 2012, Lowen et al. 2013, Robinson et al. 2014, Babarro et al. 2016). For some cases, experimental evidence indicates that such modifications increase the resistance to predation (Boulding 1984, Norberg & Tedengren 1995, Reimer & Tedengren 1996, Freeman 2007, Newell et al. 2007, Robinson et al. 2014). Mussels have been useful model systems in this regard. For example, when exposed to cues from sea stars, mussels develop thicker and more rounded shells, stronger adductor muscles, and more byssal threads (Côté 1995, Reimer & Tedengren 1997, Leonard et al. 1999, Reimer & Harms-Ringdahl 2001, Freeman & Byers 2006, Freeman 2007, Shin et al. 2009, Lowen et al.(2013). Live-feeding experiments have shown that those responses improve mussel survival and increase the time and energy required for sea stars to pry open mussels (Norberg & Tedengren 1995, Reimer & Tedengren 1996, 1997, Freeman 2007, Lowen et al. 2013). When exposed to crab cues, mussels increase shell thickness –sometimes at the expense of shell elongation– (Leonard et al. 1999, Caro & Castilla 2004, Cheung et al. 2004, Freeman & Byers 2006,Freeman 2007, Shin et al. 2009, Lowen et al. 2013, Naddafi & Rudstam 2014) and reinforce attachment to the substrate (Wang et al. 2010, Lowen et al. 2013). Shell thickening increases the handling time and force required for crabs to crush and consume a mussel (Boulding 1984, Leonard et al. 1999, Freeman 2007).

Drilling predators, such as many snail species, are also common predators of mussels worldwide. When exposed to cues from such predators, mussels also respond by thickening their shells (Smith & Jennings 2000, Cheung et al. 2004, Freeman 2007, Babarro et al. 2016). However, whether such modifications improve the ability of mussels to cope with drilling predators has not been experimentally evaluated as yet. Moreover, the studies that have shown that predator-induced morphological plasticity in bivalves hampers predation were done using lab-reared organisms, which is a less realistic approach than using organisms raised under natural conditions (Weissburg et al. 2014). To address these knowledge gaps, we conducted experiments using intertidal mussels and dogwhelks from Atlantic Canada. First, we tested the hypothesis that, in the presence of waterborne dogwhelk cues in the field, mussels would thicken their shells but grow less in length. Then, assuming the predicted shell thickening, we tested the hypothesis that the handling time required by dogwhelks to prey upon mussels would be higher when consuming mussels that were previously exposed to dogwhelk cues in the field.

## MATERIALS AND METHODS

### Effects of dogwhelk cues on mussel traits

To evaluate the effects of dogwhelk cues on mussel traits, we did a field experiment in rocky intertidal habitats from Deming Island (between 45° 12’ 41” N, 61° 10’ 50” W and 45° 12’ 45” N, 61° 10’ 26” W), Nova Scotia, Canada, during the spring and summer of 2016. The substrate of the studied habitats is stable bedrock. Maximum water velocity measured with dynamometers (Bell & Denny 1994) in these habitats was 6.0 ± 0.4 m s^-1^ (mean ± SE, n = 24), indicating that wave exposure was moderate, as values can reach 12 m s^-1^ at exposed sites in Nova Scotia (Hunt & Scheibling 2001). Intertidal temperature measured every 30 min during the study period with Hobo Pendant loggers (Onset Computer Corp., Pocasset, MA, USA) attached to the substrate was 14.2 ± 0.1 °C (n = 9 loggers). Seawater salinity measured with a handheld RF20 refractometer (Extech Instruments, Boston, MA, USA) was 35 %o. We used *Nucella lapillus*, which is the only local intertidal dogwhelk (Scrosati & Heaven 2007), and *Mytilus edulis* and *M. trossulus*, which are the two local intertidal mussels and an important prey item for dogwhelks (Largen 1967). It is highly difficult to visually identify both mussels because of morphological similarities. However, recent genetic studies have revealed that *M. trossulus* predominates over *M. edulis* in moderately exposed habitats on this coast (Tam & Scrosati 2014). Thus, given that we collected mussels at random for this study, our samples likely exhibited a predominance of *M. trossulus* over *M. edulis*.

We evaluated dogwhelk cue effects on mussel traits by manipulating dogwhelk presence in cages attached to the intertidal substrate. Each cage (Fig. 1) was made of a PVC ring (25 cm in diameter and 2.5 cm tall) and plastic mesh (0.5 cm × 0.5 cm of opening size). Each cage was divided by mesh into a central compartment (12 cm × 12 cm) and a peripheral compartment (area = 347 cm^2^). We used the peripheral compartment to create two dogwhelk treatments: either 10 enclosed dogwhelks (2.23 ± 0.02 cm long, n = 104) or none. The used dogwhelk density (ca. 3 individuals dm^-2^) was representative of the studied coast (Ellrich & Scrosati 2016). The central compartment of each cage contained a conical mesh compartment (6 cm in base diameter and 2.5 cm tall) that enclosed two mussels (1.86 ± 0.01 cm, n = 120) and, left to attach freely around the conical compartment, 18 mussels (3.5 ± 0.1 cm long, n = 30) to simulate a natural mussel patch. One of the two mussels in the conical compartment was eventually used to take the growth measurements, while the other mussel was used for the lab experiment on handling time described below. The size of the enclosed mussels is favourable for dogwhelk predation (Hughes & Dunkin 1984), which suggested that it was also suitable to detect NCEs. The cages were attached to the substrate with screws and PVC tiles, previously removing seaweeds and invertebrates to prevent potential influences (Beermann et al. 2013). Any dogwhelks found near the cages during the experiment were also periodically removed.

**Fig. 1.**
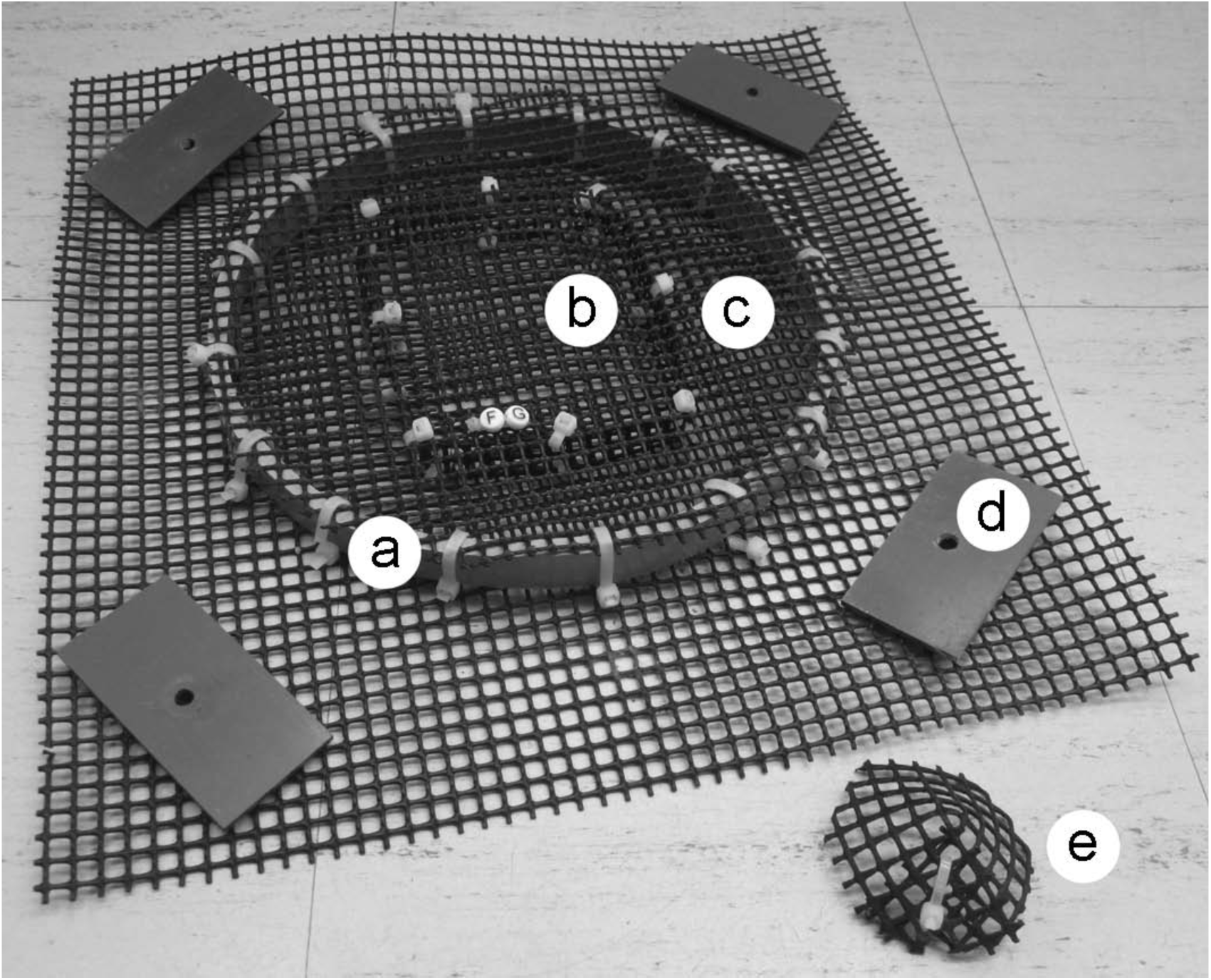
Experimental cage, showing (a) the outer PVC ring, (b) the central compartment, (c) the peripheral compartment, (d) one of the four PVC tiles used to keep the cage attached to the intertidal substrate, and (e) the conical compartment that was kept inside the central compartment holding two experimental mussels during the experiment. The PVC ring is 25 cm in diameter and 2.5 cm tall.

The experiment was arranged as a randomized complete block design with replicated treatments within blocks (Quinn & Keough 2002). We established 15 blocks at an intertidal elevation of 0.99 ± 0.04 m above chart datum, or lowest normal tide (the tidal range is 1.8 m). Each block included two replicates of each of the two dogwhelk treatments, thus yielding a total of 60 cages (30 cages per dogwhelk treatment). We started the experiment on 6 June 2016. During the experiment, we did not feed the caged dogwhelks but, to prevent their starvation, we replaced them every 10 days with mussel-fed dogwhelks kept in separate cages nearby. We used mussel-fed dogwhelks to elicit strong responses in the experimental mussels, as prey often reacts most strongly to cues from predators fed conspecific prey (Hagen et al. 2002, Schoeppner & Relyea 2005, Weissburg & Beauvais 2015). We ended the experiment on 16 August 2016, time at which we transported the mussels from the conical mesh compartments to the laboratory.

In the laboratory, we randomly selected one of the two mussels from each conical compartment for measurements. For each selected mussel, we measured shell length to the nearest 0.01 mm using a digital vernier caliper. Since we had also measured shell length at the beginning of the experiment (marking one of the two mussels in each conical compartment with nail polish for later identification), we measured relative length increment as [(*L*_*f*_ - *L_i_*)/*L*_*i*_], where *L_f_* was final length and *L_i_* was initial length. Using a vernier caliper modified with metal extensions attached to the tip of each caliper jaw, we also measured shell thickness at the lip (1 mm from the apex), centre, and base (1 mm from the base) of the right valve (looking from a dorsal view) of each mussel. Then, we dried the mussels at 50 °C for 72 hours. After that, we separated the soft tissues from the shells and measured shell mass and soft tissue dry mass to the nearest 0.1 mg.

### Effects of mussel shell thickness on dogwhelk handling time

To evaluate the effects of mussel shell thickness on dogwhelk handling time, we did a lab experiment based on the finding (see Results) that shells thickened in the presence of dogwhelk cues in the field. For the lab experiment, we used from each conical compartment the mussel that was not used for the measurements described above. Because some mussels died during the field experiment, we used 26 mussels from the dogwhelk-presence treatment and 22 mussels from the no-dogwhelk treatment. We started the lab experiment on 17 August 2016, having kept the mussels overnight since collected from the field in a culture room at 17 °C (water temperature on the studied coast in August). We placed each of the 48 mussels in a separate container with 250 ml of seawater. We then secured with epoxy glue the left valve of each mussel to a PVC tile at the bottom of each container, leaving the right valve facing upwards, exposed to predation. We selected the right valve because whelks bore the right valve more often than the left (Alexander et al. 2015). We allowed the glue to harden overnight and, then, we placed one dogwhelk (2.22 ± 0.01 cm long, n = 48) in each container. The dogwhelks had been previously starved for 10 days to standardize starvation level, which could have otherwise affected their feeding rate (Hughes & Drewett 1985, Bayne & Scullard 1978). We attached a GoPro Hero4 Black camera (GoPro, San Mateo, CA, USA) to the ceiling of the culture room to take pictures of the entire set of containers every 30 sec. We checked the containers every two hours and changed their seawater (collected on the studied coast) daily using a pipette to minimize disturbance. We ran the experiment for 18 days until 3 September 2016, although no dogwhelks fed after the thirteenth day. We measured handling time from the moment when a dogwhelk mounted its prey to the moment when the dogwhelk moved away from the formed borehole (or, in one case, when the mussel shell was empty – see Results). To confirm that shell thickening was higher in the consumed cue-exposed mussels than in the consumed mussels not exposed to cues, we measured shell thickness around the borehole of each consumed mussel. Finally, to evaluate if dogwhelks bore into a shell at points of reduced thickness, for each consumed mussel we also measured shell thickness at five intact random points on the shell.

### Data analyses

We tested the effects of dogwhelk cues (fixed factor with two levels: dogwhelk presence and absence) on shell thickness at the lip, centre, and base of mussel shells, on mussel relative length increment, on mussel shell mass, and on mussel soft tissue dry mass through separate analyses of variance (ANOVAs) appropriate for a randomized complete block design with replicated treatments within blocks (random factor with 15 levels). The assumptions of normality and homoscedasticity were tested for each variable with the Kolmogorov-Smirnov test and Cochran’s C-test, respectively (Quinn & Keough 2002). Such assumptions were met using the raw data for relative length increment, shell mass, and thickness at the centre of the shell, and using square-root-transformed data for thickness at the lip and base of the shell and soft tissue dry mass. We compared dogwhelk handling time and mussel shell thickness at the borehole between cue-exposed mussels and mussels without a previous cue exposure through independent-samples *t*-tests. Separately for each cue treatment, we compared shell thickness between the borehole area and intact shell areas (mean of the five measurements per mussel) through a paired-samples *t*-test. We did these analyses with STATISTICA 12.5 (Statsoft, Tulsa, OK, USA).

## RESULTS

At the intertidal zone, waterborne dogwhelk cues elicited an increase in the thickness of the lip, centre, and base of mussel shells but a slower growth in terms of length (Table 1, Fig. 2). Relative to the no-cue treatment, mean thickness in the presence of cues increased more (87 %) in thinner areas of the shell (lip) than in thicker areas (32 % increase at the shell centre and 47 % at the shell base; Fig. 2). Neither the mass of mussel shells nor the dry mass of mussel soft tissues was affected by dogwhelk cues (Table 1, Fig. 2). The blocks factor was only significant in one case (relative length increment, Table 1), but that result merely indicates that relative length increment varied among blocks. The important result is that the interaction between the dogwhelks factor and the blocks factor was not significant for any case, indicating that the presence or absence of NCEs, depending on the case as described above, was spatially consistent.

**Table 1.** Summary results of the analyses of variance conducted to evaluate the effects of dogwhelk cues on various mussel traits through a field experiment.

|  | SS | df | MS | *F* | p |
| --- | --- | --- | --- | --- | --- |
| **Shell thickness (lip)** |  |  |  |  |  |
| Dogwhelks | 0.696 | 1 | 0.696 | 61.56 | < 0.001 |
| Blocks | 0.173 | 14 | 0.012 | 1.09 | 0.437 |
| Dogwhelks x Blocks | 0.158 | 14 | 0.011 | 1.34 | 0.240 |
| Residual | 0.252 | 30 | 0.008 |  |  |
| **Shell thickness (centre)** |  |  |  |  |  |
| Dogwhelks | 0.267 | 1 | 0.267 | 10.11 | 0.007 |
| Blocks | 0.334 | 14 | 0.024 | 0.90 | 0.574 |
| Dogwhelks x Blocks | 0.370 | 14 | 0.026 | 1.45 | 0.190 |
| Residual | 0.545 | 30 | 0.018 |  |  |
| **Shell thickness (base)** |  |  |  |  |  |
| Dogwhelks | 0.300 | 1 | 0.300 | 21.00 | < 0.001 |
| Blocks | 0.094 | 14 | 0.007 | 0.47 | 0.915 |
| Dogwhelks x Blocks | 0.200 | 14 | 0.014 | 1.25 | 0.292 |
| Residual | 0.342 | 30 | 0.011 |  |  |
| **Relative length increment** |  |  |  |  |  |
| Dogwhelks | 0.019 | 1 | 0.019 | 18.53 | < 0.001 |
| Blocks | 0.142 | 14 | 0.010 | 9.65 | < 0.001 |
| Dogwhelks x Blocks | 0.015 | 14 | 0.001 | 0.25 | 0.996 |
| Residual | 0.128 | 30 | 0.004 |  |  |
| **Shell mass** |  |  |  |  |  |
| Dogwhelks | 0.020 | 1 | 0.021 | 1.02 | 0.329 |
| Blocks | 0.284 | 14 | 0.020 | 1.00 | 0.504 |
| Dogwhelks x Blocks | 0.285 | 14 | 0.020 | 1.00 | 0.479 |
| Residual | 0.612 | 30 | 0.020 |  |  |
| **Soft tissue dry mass** |  |  |  |  |  |
| Dogwhelks | 0.001 | 1 | 0.001 | 0.87 | 0.368 |
| Blocks | 0.014 | 14 | 0.001 | 0.91 | 0.567 |
| Dogwhelks x Blocks | 0.015 | 14 | 0.001 | 0.95 | 0.526 |
| Residual | 0.034 | 30 | 0.001 |  |  |

**Fig. 2.**
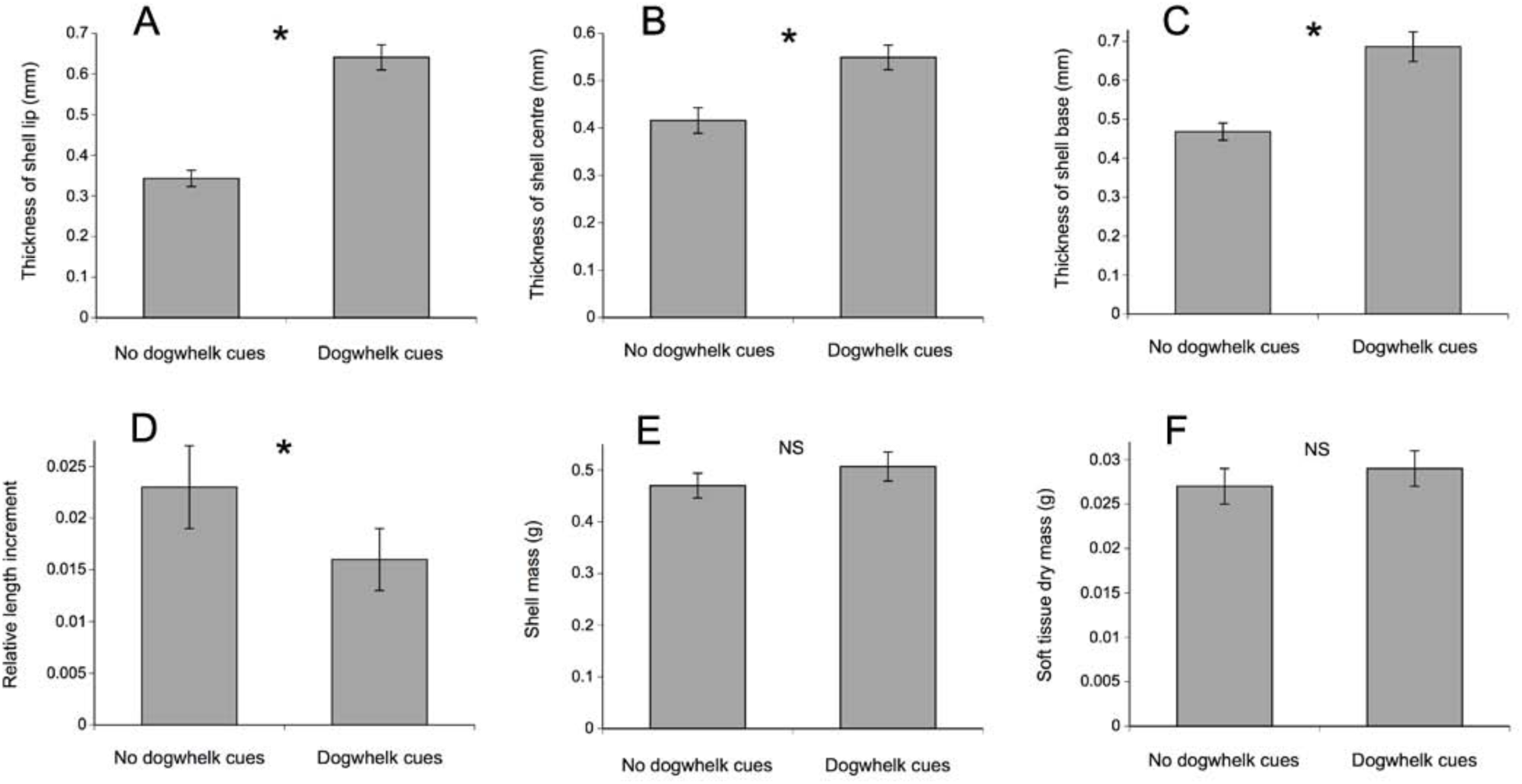
Mussel traits (mean ± SE) depending on the presence or absence of dogwhelk cues: (A) thickness at the lip, (B) centre, and (C) base of mussel shells, (D) relative length increment of shells, (E) shell mass, and (F) soft tissue dry mass. Asterisks indicate a significant difference between both treatments, whereas “NS” indicates a nonsignificant result.

In the lab experiment, 16 dogwhelks consumed their provided mussel by drilling a hole in the mussel's shell. One dogwhelk waited for its mussel to gape to then insert its proboscis through the opening, while the remaining tested dogwhelks made no apparent attempt to feed. The drilling dogwhelks needed a longer handling time to consume mussels previously exposed to dogwhelk cues compared with mussels without a cue exposure (*t*_12_ = 1.91, p = 0.040; Fig. 3). Data for two drilling dogwhelks could not be used for that t-test because such dogwhelks started to handle their respective mussel during a short initial period when the camera did not record images. Handling time was accurately calculated for all of the other dogwhelks. Shell thickness at the borehole was higher in mussels previously exposed to dogwhelk cues than in mussels without a previous cue exposure (*t*_14_ = 5.02, p < 0.001; Fig. 3). Shell thickness at the borehole was lower than at intact parts of the shell regardless of previous cue exposure (*t*_8_ = 5.50, p <0.001 for cue-exposed mussels and *t*_6_ = 4.89, p = 0.003 for mussels not previously exposed to cues; Fig. 3). Six mussels released gametes to the water soon after a dogwhelk was introduced in the container.

**Fig. 3.**
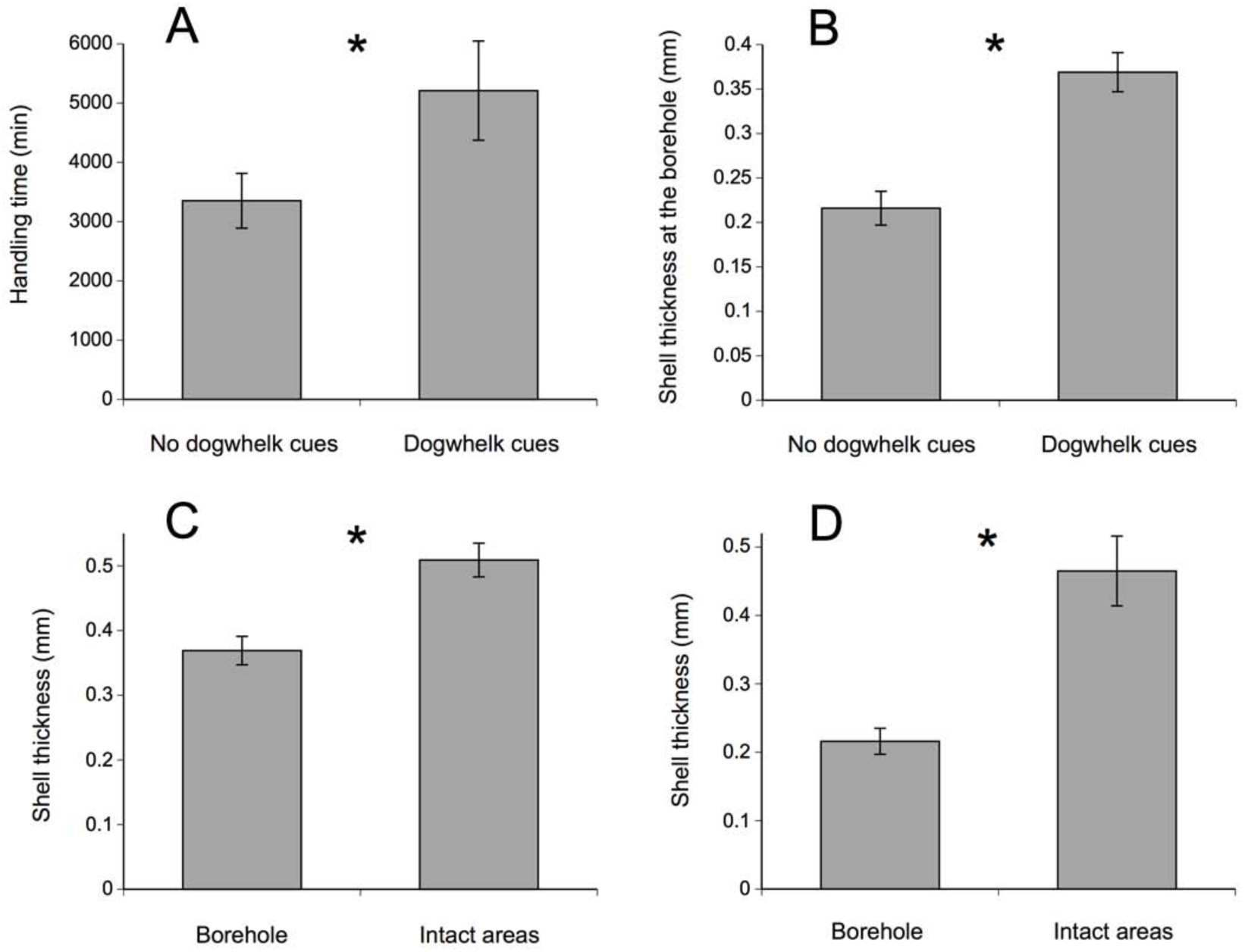
(A) Handling time by dogwhelks (mean ± SE) and (B) shell thickness at the borehole of mussels previously exposed to dogwhelk cues and mussels not exposed to such cues. Shell thickness at the borehole and at intact shell areas of mussels (C) previously exposed to dogwhelk cues and (D) mussels not exposed to such cues. Asterisks indicate a significant difference between both treatments.

## DISCUSSION

The present study has revealed that mussels in natural environments respond to dogwhelk cues by thickening their shells, although simultaneously limiting linear shell growth. Since the mass of mussel shells was not affected by dogwhelk presence, a trade-off between shell thickening and elongation was identified. Such a plastic trade-off has been observed for mussels in general in response to crab cues (Reimer & Tedengren 1997, Leonard et al. 1999, Freeman 2007, Shin et al. 2009, Lowen et al. 2013, Naddafi & Rudstrom 2014) and cues from other whelks (Smith & Jennings 2000, Freeman 2007). For other bivalves (i.e., oysters), changes in shell thickness come at a cost to soft tissue growth (Robinson et al. 2014). However, there has been no indication of such a trade-off for mussels (Reimer & Tedengren 1997, Cheung et al. 2004, Babarro et al. 2016) and the lack of dogwhelk effects on mussel soft tissue mass in our study supports that notion. Besides their NCEs on mussel shells, dogwhelks also triggered short-term responses, as some mussels exhibited a broadcast reproductive response to direct predatory threat in the lab experiment. Such a response may have been a last attempt by mussels to spawn, which aligns with findings that *Mytilus edulis* reacts to predator cues by escalating short-term investments in reproductive output (Reimer 1999).

This study also confirms that induced shell thickening in mussels occurs throughout the entire shell (Leonard et al. 1999) and it shows for the first time that thickening varies across the shell. Thickening resulting from exposure to dogwhelk cues was most pronounced in thinner areas of the shell, which are potentially most vulnerable to drilling by predators. Despite the overall thickening of mussel shells, however, dogwhelks were still able to find the thinnest part to drill, as shell thickness at the borehole was lower than at intact parts of the shell.

The functional significance of the plastic response in mussels observed in the field was revealed by the lab experiment as, on average, dogwhelks took 55 % longer to handle to full consumption mussels that had been exposed to dogwhelk cues and, hence, had thicker shells. These results show for the first time that predator-induced shell thickening in mussels hinders the feeding of drilling predators. Induced morphological defenses in mussels also increase handling time for predatory crabs (Boulding 1984, Reimer & Tedengren 1997, Freeman 2007) and sea stars (Norberg & Tedengren 1995, Freeman 2007). However, comparable proportional increases in handling time may be more detrimental for dogwhelks. The regular handling time required for dogwhelks to drill into mussels is considerably longer than for crabs and sea stars to crush or pry open a mussel (Freeman 2007, Miller 2013). Thus, a similar proportional increase in handling time would expose dogwhelks considerably longer than those predators to desiccation stress (Hughes & Dunkin 1984, Davenport & MacAlister 1996), to competitors that can displace a feeding dogwhelk (Hughes & Dunkin 1984, Quinn et al. 2012, Chattopadhyay & Baumiller 2007, Hutchings & Herbert 2013), to predators (Vadas et al. 1994), to entrapment by neighboring mussels (Davenport et al. 1996, Farrell & Crowe 2007), and to dislodgement by waves (Denny 1988). Thus, the morphological alterations that increase the time a mussel is handled by a dogwhelk should increase the likelihood that the mussel survives a predation attempt.

Future research could investigate if the dogwhelk-induced responses differ to some extent between *Mytilus edulis* and *M. trossulus*. Recent research has found that *M. edulis* exhibits stronger morphological responses to crab and sea star cues than *M. trossulus* (Lowen et al. 2013), but this may not be the case when presented with more reliable cues from slower-moving predators such as dogwhelks (Burrows & Hughes 1991). Since cue release by dogwhelks increases their vulnerability to byssal entrapment by mussels (Farrell & Crowe 2007) and elicits prey structural changes that reduce feeding success, future studies could also investigate behavioural adaptations of dogwhelks to begin feeding during low tides to delay cue dissemination in the water.

Overall, this is the first study that has used organisms raised in the field, rather than in the lab, to demonstrate that predator-induced morphological responses in bivalve prey hinder predation. The complex abiotic and biotic conditions of intertidal environments, almost impossible to replicate in a lab, conferred a high degree of realism to the results from the field experiment and the organisms used for the feeding assay. This approach is thus in line with recent studies that have highlighted the need to understand NCEs under realistic conditions in order to improve theory development (Weissburg et al. 2014, Babarro et al. 2016).

## Acknowledgements

We thank Sonja Ehlers and Jadine Krist for field assistance. This project was funded by grants from the Natural Sciences and Engineering Research Council of Canada (NSERC Discovery Grant) and the Canada Foundation for Innovation (CFI) to RAS.

## LITERATURE CITED

Alexander ME, Raven HJ, Robinson TB (2015) Foraging decisions of a native whelk, *Trochia cingulata* Linnaeus, and the effects of invasive mussels on prey choice. J Exp Mar Biol Ecol 470:26–33

Babarro JMF, Vázquez E, Olabarria C (2016) Importance of phenotypic plastic traits on invasive success: response of *Xenostrobus securis* to the predatory dogwhelk *Nucella lapillus*.Mar Ecol Prog Ser 560:185–198

Bayne BL, Scullard C (1978) Rates of feeding by *Thais (Nucella) lapillus* (L.). J Exp Mar Biol Ecol 32:113–129

Beermann AJ, Ellrich JA, Molis M, Scrosati RA (2013) Effects of seaweed canopies and adult barnacles on barnacle recruitment: the interplay of positive and negative influences. J Exp Mar Biol Ecol 448:162–170

Bell EC, Denny MW (1994) Quantifying “wave exposure”: a simple device for recording maximum velocity and results of its use at several sites. J Exp Mar Biol Ecol 156:199–215

Boulding EG (1984) Crab-resistant features of shells of burrowing bivalves: decreasing vulnerability by increasing handling time. J Exp Mar Biol Ecol 76:201–223

Brönmark C, Hansson LA (2012) Chemical ecology in aquatic systems. Oxford University Press, Oxford

Burrows MT, Hughes RN (1991) Variation in foraging behavior among individuals and populations of dogwhelks, *Nucella lapillus*: natural constraints on energy intake. J Anim Ecol 60:497–514

Caro AU, Castilla JC (2004) Predator-inducible defenses and local intra-population variability of the intertidal mussel *Semimytilus algosusin* in central Chile. Mar Ecol Prog Ser 276:115–123

Chattopadhyay D, Baumiller TK (2007) Drilling under threat: an experimental assessment of the drilling behavior of *Nucella lamellosa* in the presence of a predator. J Exp Mar Biol Ecol 352:257–266

Cheung SG, Lam S, Gao QF, Mak KK, Shin PKS (2004) Induced anti-predator responses of the green mussel, *Perna viridis* (L.), on exposure to the predatory gastropod, *Thais clavigera* Kuster, and the swimming crab, Thalamita danae Stimpson. Mar Biol 144:675–684

Côté IM (1995) Effects of predatory crab effluent on byssus production in mussels. J Exp Mar Biol Ecol 188:233–241

Davenport J, MacAlister H (1996) Environmental conditions and physiological tolerances of intertidal fauna in relation to shore zonation at Husvik, South Georgia. J Mar Biol Assoc U K 76:985–1002

Davenport J, Moore PG, LeComte E (1996) Observations on defensive interactions between predatory dogwhelks, *Nucella lapillus* (L.), and mussels, *Mytilus edulis* L. J Exp Mar Biol Ecol 206:133–147

Denny MW (1988) Biology and the mechanics of the wave swept environment. Princeton University Press, Princeton

Ellrich JA, Scrosati RA, Molis M (2015) Predator nonconsumptive effects on prey recruitment weaken with recruit density. Ecology 96:611–616

Ellrich JA, Scrosati RA (2016) Water motion modulates predator nonconsumptive limitation of prey recruitment. Ecosphere 7:e01402

Farrell ED, Crowe TP (2007) The use of byssus threads by *Mytilus edulis* as an active defense against Nucella lapillus. J Mar Biol Assoc U K 87:559–564

Ferrari MCO, Wisenden BD, Chivers DP (2010) Chemical ecology of predator–prey interactions in aquatic ecosystems: a review and prospectus. Can J Zool 88:698–724

Freeman AS (2007) Specificity of induced defenses in *Mytilus edulis* and asymmetrical predator deterrence. Mar Ecol Prog Ser 334:145–153

Freeman AS, Byers JE (2006) Divergent induced responses to an invasive predator in marine mussel populations. Science 313:831–833

Hagen NT, Andersen A, Stabell OB (2002) Alarm responses of green sea urchin, *Strongylocentrotus droebachiensis*, induced by chemically labeled durophagous predators and simulated acts of predation. Mar Biol 140:365–374

Hughes RN, Drewett D (1985) A comparison of foraging behavior of dogwhelks, *Nucella lapillus* (L.), feeding on barnacles or mussels on the shore. J Mollusc Stud 51:73–77

Hughes RN, Dunkin S (1984) Behavioral components of prey selection by dogwhelks, Nucella lapillus (L.), feeding on mussels, *Mytilus edulis* L., in the laboratory. J Exp Mar Biol Ecol 77:45–68

Hunt HL, Scheibling RE (2001) Patch dynamics of mussels on rocky shores: integrating process to understand pattern. Ecology 82:3213–3231

Hutchings JA, Herbert GS (2013) No honor among snails: conspecific competition leads to incomplete drill holes by a naticid gastropod. Palaeogeogr Palaeoclimatol Palaeoecol 379–380:32–38

Keppel E, Scrosati R (2004) Chemically mediated avoidance of *Hemigrapsus nudus* (Crustacea) by Littorina scutulata (Gastropoda): effects of species coexistence and variable cues. Anim Behav 68:915–920

Largen MJ (1967) The diet of the dog-whelk, *Nucella lapillus* (Gastropoda Prosobranchia). J Zool 151:123–127

Leonard GH, Bertness MD, Yund PO (1999) Crab predation, waterborne cues, and inducible defenses in the blue mussel, Mytilus edulis. Ecology 80:1–14

Lord JP, Whitlatch RB (2012) Inducible defenses in the eastern oyster *Crassostrea virginica* Gmelin in response to the presence of the predatory oyster drill *Urosalpinx cinerea* Say in Long Island Sound. Mar Biol 159:1177–1182

Lowen JB, Innes DJ, Thompson RJ (2013) Predator-induced defenses differ between sympatric *Mytilus edulis* and *M. trossulus*. Mar Ecol Prog Ser 475:135–143

Matassa CM, Donelan SC, Luttbeg B, Trussell GC (2016) Resource levels and prey state influence antipredator behavior and the strength of nonconsumptive predator effects. Oikos 125:1478–1488

Miller LP (2013) The effect of water temperature on drilling and ingestion rates of the dogwhelk Nucella lapillus feeding on Mytilus edulis mussels in the laboratory. Mar Biol 160:1489–1496

Molis M, Preuss I, Firmenich A, Ellrich J (2011) Predation risk indirectly enhances survival of seaweed recruits but not intraspecific competition in an intermediate herbivore species. J Ecol 99:807–817

Naddafi R, Rudstam LG (2014) Predator-induced morphological defenses in two invasive dreissenid mussels: implications for species replacement. Freshwat Biol 59:703–713

Nakaoka M (2000) Nonlethal effects of predators on prey populations: predator-mediated change in bivalve growth. Ecology 81:1031–1045

Newell RIE, Kennedy VS, Shaw KS (2007) Comparative vulnerability to predators, and induced defense responses, of eastern oysters *Crassostrea virginica* and non-native *Crassostrea ariakensis oysters* in Chesapeake Bay. Mar Biol 152:449–460

Norberg J, Tedengren M (1995) Attack behavior and predatory success of *Asterias rubens* L. related to differences in size and morphology of the mussel *Mytilus edulis* L. J Exp Mar Biol Ecol 186:207–220

Quinn BK, Boudreau MR, Hamilton DJ (2012) Inter- and intraspecific interactions among green crabs (*Carcinus maenas*) and whelks (*Nucella lapillus*) foraging on blue mussels (*Mytilus edulis*). J Exp Mar Biol Ecol 412:117–125

Quinn GP, Keough MJ (2002) Experimental design and data analysis for biologists. Cambridge University Press, Cambridge

Reimer O (1999) Increased gonad ratio in blue mussel, *Mytilus edulis*, exposed to starfish predators. Aquat Ecol 33:185–192

Reimer O, Harms-Ringdahl S (2001) Predator-inducible changes in blues mussels from the predator-free Baltic Sea. Mar Biol 139:959–965

Reimer O, Tedengren M (1996) Phenotypical improvements of morphological defenses in the mussel *Mytilus edulis* induced by exposure to the predator *Asterias rubens*. Oikos 75:383–390

Reimer O, Tedengren M (1997) Predator-induced changes in byssal attachment, aggregation, and migration in the blue mussel, *Mytilus edulis*. Mar Fresh Behav Physiol 30:251–266

Robinson EM, Lunt J, Marshall CD, Smee DL (2014) Eastern oysters *Crassostrea virginica* deter crab predators by altering their morphology in response to crab cues. Aquat Biol 20:111–118

Schoeppner NM, Relyea RA (2005) Damage, digestion, and defense: the roles of alarm cues and kairomones for inducing prey defenses. Ecol Lett 8:505–512

Scrosati R, Heaven C (2007) Spatial trends in community richness, diversity, and evenness across rocky intertidal environmental stress gradients in eastern Canada. Mar Ecol Prog Ser 342:1–14

Shin PKS, Yang FY, Chiu MY, Cheung SG (2009) Cues from the predator crab *Thalamita danae* fed different prey can affect scope for growth in the prey mussel *Perna viridis*. Mar Fresh Behav Physiol 42:343–355

Smith LD, Jennings JA (2000) Induced defensive responses by the bivalve *Mytilus edulis* to predators with different attack modes. Mar Biol 136:461–469

Tam JC, Scrosati RA (2014) Distribution of cryptic mussel species (*Mytilus edulis* and *M. trossulus*) along wave exposure gradients on northwest Atlantic rocky shores. Mar Biol Res 10:51–60

Vadas RL, Burrows MT, Hughes RN (1994) Foraging strategies of dogwhelks, *Nucella lapillus* (L.): interacting effects of age, diet, and chemical cues to the threat of predation. Oecologia 100:439–450

Wang Y, Hu M, Shin PKS, Cheung SG (2010) Induction of anti-predator responses in green lipped mussel *Perna viridi* sunder hypoxia. Mar Biol 157:747–754

Weissburg M, Beauvais J (2015) The smell of success: the amount of prey consumed by predators determines the strength and range of cascading non-consumptive effects. PeerJ 3:e1426

Weissburg M, Smee DL, Ferner MC (2014) The sensory ecology of nonconsumptive predator effects. Am Nat 184:141–157

